# Origination of the circadian clock system in stem cells regulates cell differentiation

**DOI:** 10.1101/710590

**Authors:** Kotaro Torii, Keisuke Inoue, Keita Bekki, Kazuya Haraguchi, Minoru Kubo, Yuki Kondo, Takamasa Suzuki, Hanako Shimizu, Kyohei Uemoto, Masato Saito, Hiroo Fukuda, Takashi Araki, Motomu Endo

## Abstract

The circadian clock regulates various physiological responses. To achieve this, both animals and plants have distinct circadian clocks in each tissue that are optimized for that tissue’s respective functions. However, if and how the tissue-specific circadian clocks are involved in specification of cell types remains unclear. Here, by implementing a single-cell transcriptome with a new analytics pipeline, we have reconstructed an actual time-series of the cell differentiation process at single-cell resolution, and discovered that the *Arabidopsis* circadian clock is involved in the process of cell differentiation through transcription factor BRI1-EMS SUPPRESSOR 1 (BES1) signaling. In this pathway, direct repression of *LATE ELONGATED HYPOCOTYL* (*LHY*) expression by BES1 triggers reconstruction of the circadian clock in stem cells. The reconstructed circadian clock regulates cell differentiation through fine-tuning of key factors for epigenetic modification, cell-fate determination, and the cell cycle. Thus, the establishment of circadian systems precedes cell differentiation and specifies cell types.

---

The circadian clock is involved in various physiological responses to regulate a large set of genes in both animals and plants^1^. Circadian clocks in each tissue regulate different responses consistent with tissue-specific sets of circadian-regulated genes^2,3^. The tissue-specific clocks are considered to be optimized for each tissue’s respective functions, and therefore there is a possibility that the different clock functions in each tissue contribute to specify the cell type.

In mammalian embryonic stem cells, circadian rhythms are not observed during early developmental stages, but they emerge along with cell differentiation; and the circadian rhythms in differentiated cells disappear when they are reprogrammed^4^. Consistently, several mutations in clock genes cause abnormal cell differentiation in mammals^5^. In *Arabidopsis*, a *circadian clock associated 1*; *late elongated hypocotyl* double mutant (*cca1 lhy*) shows an increased number of free-ending vascular bundles^6^, indicating abnormal vascular cell differentiation in the clock mutant.

To investigate the involvement of the plant circadian clock in cell differentiation, we performed detailed observations of clock mutants, *cca1*; *lhy*; *timing of CAB expression 1* (*cca1 lhy toc1*) as well as a knockdown of *BROTHER OF LUX ARRHYTHMO* (also known as *NOX*) by artificial microRNA in a *lux arrhythmo*/*phytoclock1* mutant background (*lux nox*). We confirmed that the clock mutants affect development of vascular bundles, guard cells, and root cells (Supplementary Fig. 1a-c), suggesting that the plant circadian clock is generally involved in the process of cell differentiation. We then utilized the vascular cell differentiation induction system, referred to as VISUAL^7^, for further investigation of the molecular mechanisms driving the clock-mediated cell differentiation. In this system, vascular cells (including both xylem and phloem cells) are induced from mesophyll cells through the stem cells and vascular stem cells (Fig. 1a). A loss-of-function mutation in *BRI1-EMS SUPPRESSOR 1* (*BES1*) showed severe defects in vascular cell differentiation, as shown previously^8^. We found that clock mutants showed significant defects in vascular cell differentiation (Fig. 1b). Perturbation of endogenous circadian rhythms by random light/dark conditions also inhibited vascular cell differentiation (Fig. 1c), suggesting the requirement of a functional circadian clock for cell differentiation.

**Figure 1.**
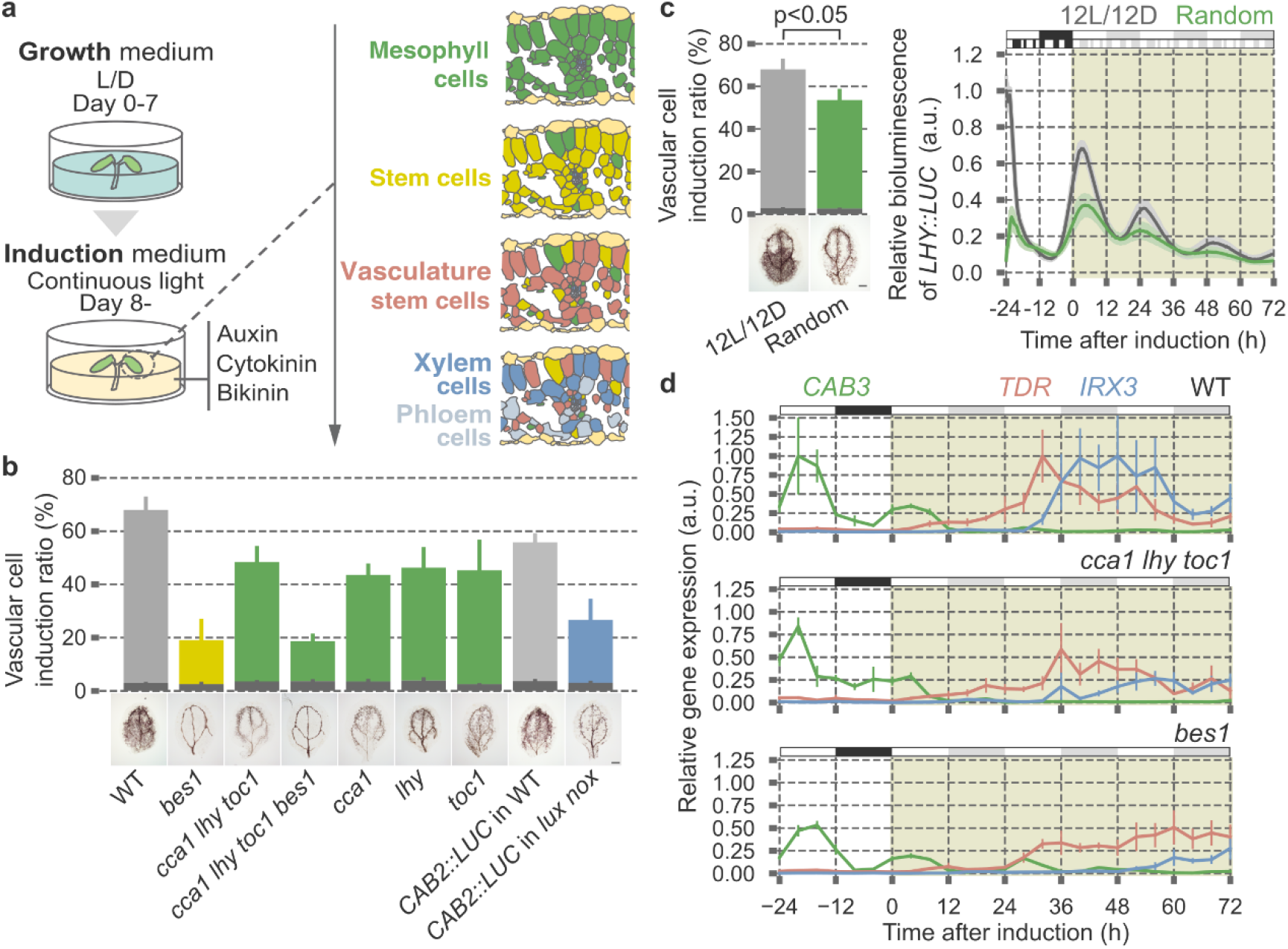
The plant circadian clock affects cell differentiation. **a**, Schematic of ectopic vascular cell induction with VISUAL. **b**, Lignin stained xylem cell density in WT, *bes1*, and clock mutants after induction. Light gray, green, and yellow bars indicate ectopically induced xylem cells. Dark gray bars indicate endogenous xylem cells (n = 5, mean ±s.e.). Representative photos of lignin staining are below (bar = 1 mm). **c**, Perturbation of endogenous circadian rhythms by random light/dark periods (2L3D1L1D3L1D1L2D3L2D2L3D, total 12L12D per 24 h) inhibits vascular cell differentiation with VISUAL. Left, lignin stained xylem cell density and representative photos (n = 5, mean ± s.e., bar = 1 mm, two-sided student’s t-test, p < 0.05). Right, expression patterns of *LHY∷LUC* during VISUAL under normal and random light/dark conditions (n = 10, mean ± s.e.). **d**, Expression patterns of cell-type-specific markers during VISUAL in WT, *cca1 lhy toc1*, and *bes1* (n = 3, mean ± s.e.). Green, red, and blue lines indicate marker genes for mesophyll cells (*CAB3*), vascular stem cells (*TDR*), and xylem cells (*IRX3*), respectively.Expression peaks of each respective gene in WT were normalized to 1. White, black, and gray boxes indicate light period, night period, and subjective night period, respectively.

The VISUAL assay consists of two steps, dedifferentiation of mesophyll cells to stem cells and differentiation of stem cells into vascular cells via vascular stem cells. To determine which step is regulated by the circadian clock, we measured expression levels of cell-type-specific markers. In the wild type (WT), a mesophyll cell marker (*CAB3*) rapidly disappeared within 24 h after induction. Afterward, a vascular stem cell marker (*TDR*) and a vascular cell marker (*IRX3*) appeared in succession (Fig. 1d and Supplementary Fig. 1d-f). *TDR* was not fully induced in *cca1 lhy toc1* and *bes1* mutants, although the reduction of *CAB3* expression in both mutants was comparable to WT (Fig. 1d). These data suggest that both the clock genes and BES1 are involved in the step of differentiation rather than dedifferentiation.

To clarify the relationship between the clock genes and BES1, we investigated a possible link between both factors. Consistent with a previous report^9^, chromatin immunoprecipitation (ChIP) using plants expressing BES1-GFP demonstrated enrichment of BES1-GFP at the G-box and E-box motifs in the *LHY* promoter (Fig. 2a). Transient co-expression of BES1 and *LHY∷LUC* resulted in decreased luciferase activity (Fig. 2b), suggesting that BES1 acts as a repressor of *LHY* expression. In our VISUAL conditions, an active dephosphorylated BES1 accumulated immediately after induction (Fig. 2c). Consistent with the accumulation of dephosphorylated BES1, the amplitude of *LHY* and *CCA1* expression decreased after induction (Fig. 2d). We next tested the effect of BES1 on *LHY* expression during VISUAL. Rhythmic *LHY* expression with low amplitude was sustained in WT, whereas the *bes1* mutation caused complete loss of *LHY* expression after induction (Fig. 2e). Given that stem cells should be enriched in *bes1* mutant after induction (Fig. 1d), the requirement of BES1 for *LHY* expression suggested that BES1 is essential for triggering circadian rhythms in the stem cells.

**Figure 2.**
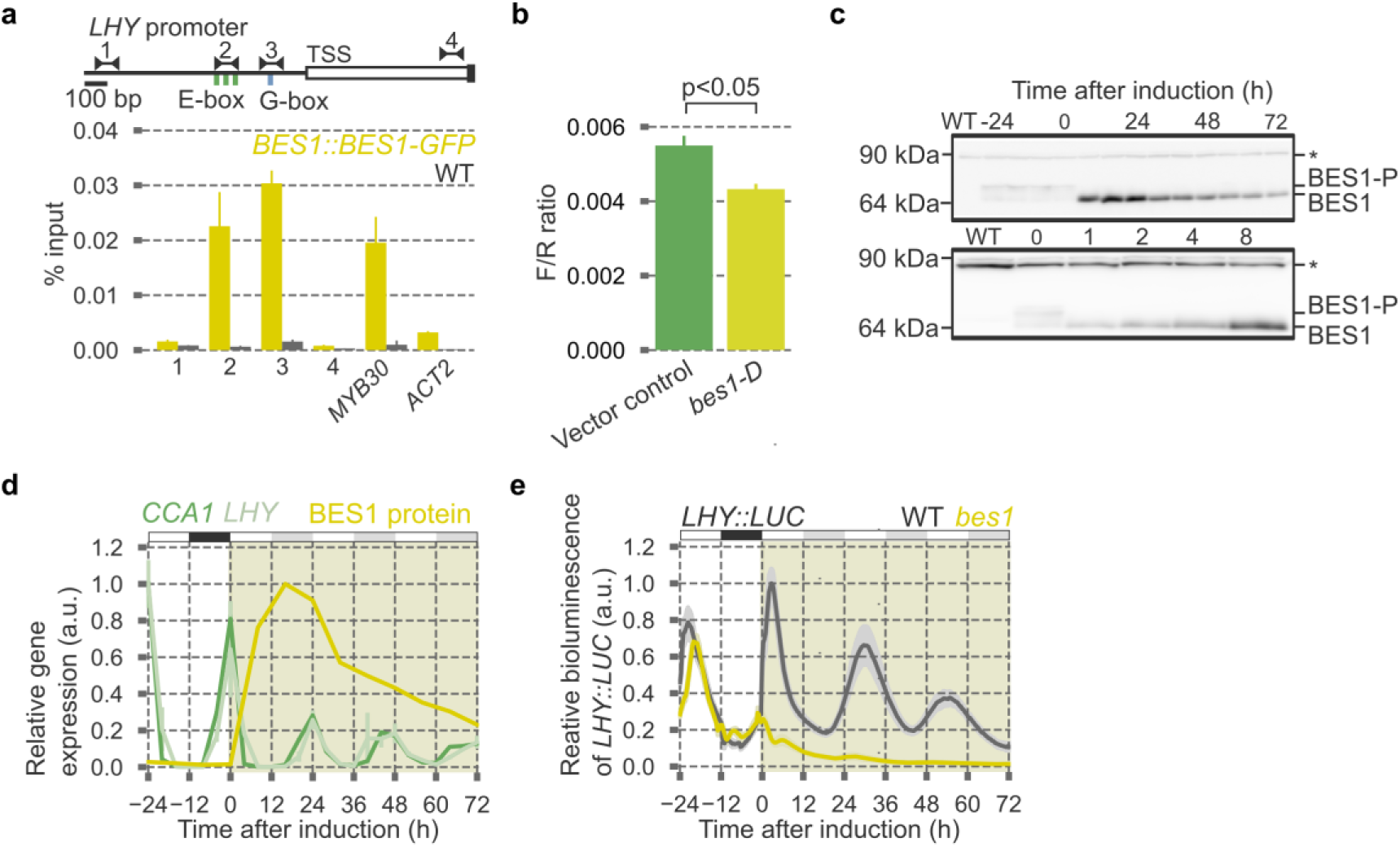
BES1 is required for *LHY* expression in stem cells. **a**, ChIP-qPCR analyses using WT and *BES1∷BES1-GFP* (n = 3, mean ± s.e.). *MYB30* and *ACT2* were used as positive and negative controls, respectively. Green and blue boxes indicate E-box (CANNTG) and G-box (CACGTG) motifs. TSS: transcriptional start site. **b**, Expression of *LHY∷LUC* using *35S∷bes1-D* as an effector in *N. benthamiana* (n = 5, mean ± s.e., two-sided student’s t-test, p < 0.05). *35S∷RLUC* was used as a transformation control. F/R ratio indicates ratio of firefly and renilla luciferase activities. **c**, Immunoblot analyses of BES1-GFP proteins during VISUAL. Samples were harvested every 8 h, from 24 h before induction up to 72 h after induction (top) and from 1 to 8 h after induction (bottom). BES1-P indicates phosphorylated BES1. Asterisks indicate non-specific bands. **d**, Expression patterns of *LHY* and *CCA1* during VISUAL (n = 3, mean ± s.e.). Relative dephosphorylated BES1 levels determined from Fig. 2c are overlaid in yellow. **e**, Expression patterns of *LHY∷LUC* in WT and *bes1* during VISUAL (n = 15, mean ± s.e.). White, black, and gray boxes indicate light period, night period, and subjective night period, respectively. An expression peak in WT was normalized to 1.

Since the expression of *TDR* and *IRX3* overlapped each other, and cell differentiation proceeded gradually, even in the VISUAL assay (Fig. 1d and Supplementary Fig. 1d-f), it was still unclear whether the development of circadian rhythms triggers cell differentiation or *vice versa*. To circumvent such low spatiotemporal resolution of bulk analysis, which can be attributed to the crude averaging of various cell types, we performed time-series single-cell RNA sequencing (scRNA-seq) with VISUAL. Although making protoplasts is the conventional way to obtain single plant cells, the procedure is not only time-consuming but also potentially stress-inducing. We therefore obtained total RNA from single cells using glass capillaries^10^ (Supplementary Fig. 2a and Supplementary Video 1). 216 single cell samples harvested every 4 h, from 24 h before induction up to 84 h after induction were subjected to RNA-seq, and the collected data were normalized together with time-series cell-population RNA sequencing (cpRNA-seq) data obtained from whole cotyledons (Supplementary Fig. 2b-e and Supplementary Table 1).

To separate xylem and phloem cell lineages, we first applied the Wishbone algorithm, which can order scRNA-seq data with bifurcating developmental trajectories^11^. t-distributed Stochastic Neighbor Embedding (t-SNE) of the dataset was represented by a Y-shaped structure, suggesting that the Wishbone can properly reconstruct developmental trajectories of xylem and phloem cells (Fig. 3a). This assertion was further validated by overlaying expression levels of cell-type-specific markers (Fig. 3b and Supplementary Fig. 3a). For simplicity, we focused on the xylem cell lineage and applied the Seurat algorithm^12^ for improving temporal resolution of the pseudo-time trajectory. Seurat allowed us to estimate the most likely location of cells by referring to preset expression patterns of cell-type-specific markers. Clustering of the data obtained by applying 24 reference genes in Seurat yielded four major clusters presumed to be mesophyll cells, stem cells, vascular stem cells, and xylem cells (Supplementary Fig. 3b and Supplementary Table 2). The stem cell state was represented by high expression of *OBP1* and *ACS6* (Fig. 3b and Supplementary Fig. 3b). *OBP1* is highly expressed in tissue with high cell proliferation activity such as developing embryo and organ primordia^13^, and *ACS6* encodes the rate-limiting enzyme for biosynthesis of ethylene, a plant hormone that promotes stem cell division in the root^14^, demonstrating the utility of the two genes as stem cell markers.

**Figure 3.**
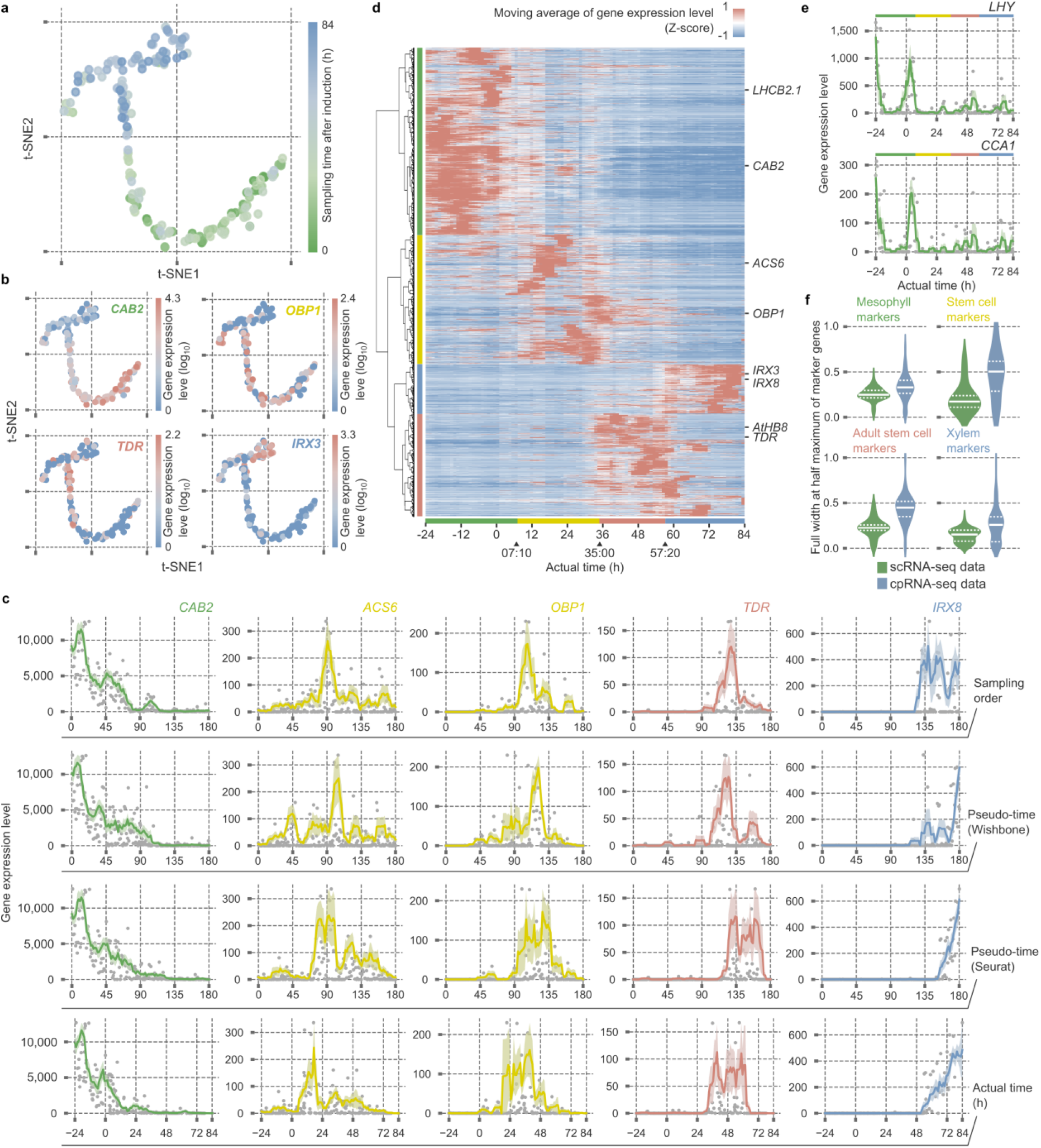
WISP pipeline improves temporal resolution of scRNA-seq data. **a**, **b**, Bifurcated cell lineage on t-SNE 2D plots predicted by Wishbone, showing sampling times, and overlaid by expression levels of cell-type-specific markers. Color codes indicate sampling times from early (green) to late (blue) (**a**) and normalized UMI counts from low (blue) to high (red) (**b**). **c**, Comparison of marker gene expression patterns aligned by sampling order, pseudo time (Wishbone), pseudo time (Wishbone-Seurat), and actual time (WISP). **d**, A hierarchically-clustered heat map visualizing Z-scores of moving averages of gene expression levels with a window size of 4 h. Green, yellow, red, and blue vertical bars indicate marker genes for mesophyll cells, stem cells, vascular stem cells, and xylem cells, respectively. **e**, Reconstruction of 24 h periodicity in time-series scRNA-seq by the WISP pipeline. Green, yellow, red, and blue bars indicate mesophyll cell, stem cell, vascular stem cell, and xylem cell states, respectively. **f**, Comparison of full width at half maximum of cell-type-specific marker expression peaks between the actual time-series scRNA-seq data and the cpRNA-seq data. White solid lines indicate the median. White broken lines indicate the upper and lower quartiles.

In general, scRNA-seq analysis kills target cells in the process of obtaining transcriptome data; thus, time-series analysis is still a daunting problem^15^. Due to the lack of temporal information, circadian rhythms have not been rigorously studied using single-cell transcriptomes. To overcome this limitation and to reconstruct actual time-series from single-cell transcriptome datasets, we developed the PeakMatch algorithm. The basic concept of PeakMatch is that the timing of significant gene expression peaks can be comparable between scRNA-seq data based on pseudo time-series and cpRNA-seq data based on actual time-series (Supplementary Fig. 3c, see Methods for details). By integrating estimated peak times of 2,217 genes, we reconstructed an actual time-series and succeeded in improving segregation of each cell-type-specific marker (Fig. 3c,d), and detected 24 h rhythmicity of *LHY* and *CCA1* expression, even in scRNA-seq data (Fig. 3e and Supplementary Table 3). The half-value width of the gene expression peaks in the reconstructed actual time-series was narrow compared to that of cpRNA-seq (Fig. 3f), demonstrating that PeakMatch reduces the crude averaging effect and improves temporal resolution. Taken together, the Wishbone-Seurat-PeakMatch (WISP) pipeline improved spatiotemporal resolution of scRNA-seq data and enabled us to handle actual time-series data at single-cell resolution.

Using this reconstructed scRNA-seq data, we evaluated the expression of clock genes during cell differentiation. Consistent with our previous report^3^, expression of *PRR5* and *PRR7*, two clock genes whose expression is predominant in mesophyll cells, was diminished in concert with the loss of mesophyll cell identity, whereas *ELF4* and *LUX*, components of Evening Complex (EC) whose expression is enriched in vascular cells, began to emerge approximately 24 h after induction (Fig. 4a,b). We also found disruptions of circadian rhythms in the stem cells, as evidenced by the expression patterns of clock genes (Supplementary Fig. 3d). Since re-phasing of circadian rhythms was also observed in root stem cell niches^16,17^, reconstruction of the circadian clock system could be generally required for cell differentiation. To test how the reconstructed clock regulates cell differentiation, we calculated the kinetics of GO-term enrichment. GO-terms related to DNA methylation and cell cycle were significantly enriched soon after the induction of *ELF4* and *LUX* (Fig. 4c). We then performed ChIP-seq analysis using plants expressing LUX-GFP to identify direct targets of the circadian clock (Supplementary Table 4). We found that *CYCD3;1*, *RBR*, and *E2Fc*, which play leading roles in the G1-S transition, were significantly enriched (Fig. 4d and Supplementary Fig. 4a). Consistently, clock mutants showed disruption of daily rhythms in cell proliferation and decreased numbers of meristematic cells, and altered cell-cycle-related gene expression and DNA ploidy patterns (Supplementary Fig. 4b-e). RBR also controls epigenetic regulation through DNA methylation during cell differentiation^18^, and E2Fc is reported as a key regulator for xylem differentiation^19^. Taken together, we concluded that the reconstructed circadian clock integratively regulates cell differentiation through fine-tuning of key factors for epigenetic modifications, cell-fate determination, and cell cycle (Fig. 4e). Our finding that BES1-triggered reconstruction of the circadian clock regulates genes related to cell cycle was further supported by re-analysis of recent scRNA-seq data^20^ derived from root tips (Supplementary Fig. 4f).

**Figure 4.**
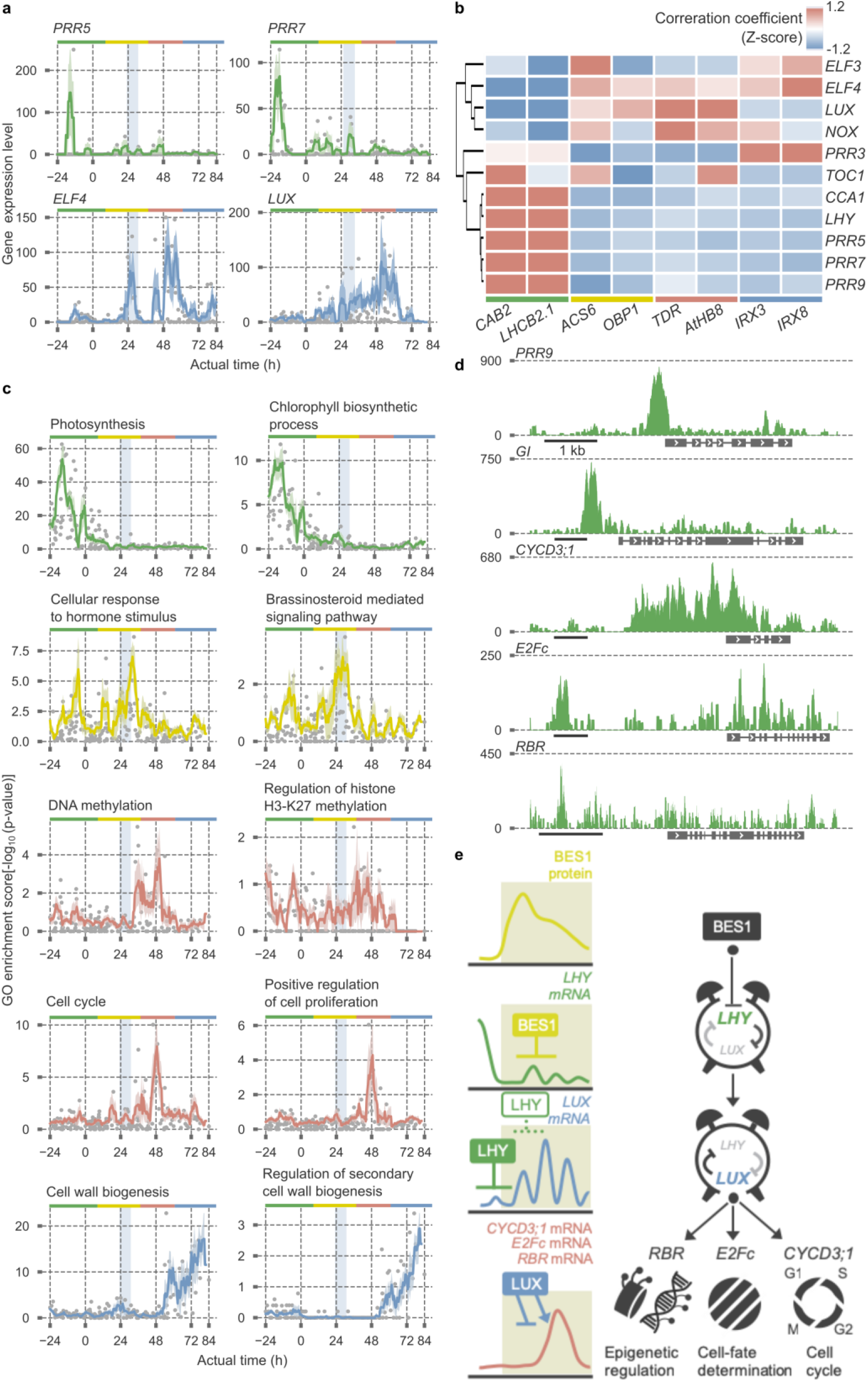
Reconstruction of circadian rhythms prior to cell-fate determination. **a**, Expression patterns of clock genes in actual time-series scRNA-seq data. Expression of *ELF4* and *LUX* originates in the stem cell. The first peak time of *ELF4* and *LUX* expression is highlighted by the pale blue window. **b**, Correlation coefficient between clock genes and cell-type-specific markers. **c**, GO-term enrichment during cell-fate transition. Genes related to cell cycle are enriched soon after the first peak of *ELF4* and *LUX* expression. The first peak time of *ELF4* and *LUX* expression is highlighted by the pale blue window. Green, yellow, red, and blue bars indicate mesophyll cell, stem cell, vascular stem cell, and xylem cell states, respectively. **d**, Visualization of ChIP-seq data around genes related to clock, cell cycle, cell-fate determination, and epigenetic regulation (bar = 1 kb). Peak counts of reads are shown. **e**, Our model proposes that BES1 represses *LHY* expression in stem cells to reconstruct the circadian clock, resulting in induction of *LUX* expression. LUX in the reconstructed clock modulates key factors for epigenetic regulation, cell-fate determination, and cell cycle, thereby inducing cell differentiation.

We have demonstrated that establishment of circadian systems precedes cell differentiation, supporting the hypothesis that construction of the circadian clock for tissue-specific functions can specify cell types. Development of circadian rhythms during differentiation, and distinct functions of circadian clocks in each tissue, are common features across kingdoms^2-4^. Our findings and the WISP pipeline provide a new avenue for further studies of circadian clocks in cell differentiation.

## Methods

### Plant material and growth conditions

All wild type and transgenic lines used here were *Arabidopsis thaliana* ecotype Columbia-0 (Col-0). Seeds were surface-sterilized and sown on 0.8% agar plates containing Murashige and Skoog medium with 0.5% sucrose or liquid media as described previously^7^. Plants were grown under L/D (12 h light and 12 h dark, 84 μmol m^−2^ s^−1^) conditions at 22°C. For induction of ectopic vascular cell differentiation, plants were entrained by L/D conditions for 7 days, although the original protocols^7^ called for growth of plants under continuous light (LL) conditions. Bikinin, 2,4-dichlorophenoxyacetic acid (2,4-D), and kinetin were added to the eight-day-old plants at ZT0. *cca1-1 lhy-11 toc1-2* and *cca1-1 lhy-11* were provided by Takafumi Yamashino^21^ (Nagoya University), *lux-4 nox* and its parental *CAB2∷LUC* were provided by Dmitri A. Nusinow^22^ (Donald Danforth Plant Science Center), *LUX∷LUX-GFP*/*lux-4* was provided by Philip A. Wigge^23^ (University of Cambridge).

### Measurement of cell differentiation, cell proliferation, and cell cycles

For lignin staining, plants were fixed in acetic ethanol fixative (75% glacial acetic acid and 25% ethanol) for 1 day, stained with 20% phloroglucinol in 99.5% ethanol and concentrated HCl (1:19, v/v) for 1 h, cleared with chloral hydrate/glycerol/H2O mixture (8 g of chloral hydrate in 1 mL of glycerol and 2 mL of H2O) for 2 h, and observed under a light microscope. For quantification of the vascular cell induction ratio, areas for xylem cells in the cotyledons were calculated using ImageJ.

For measurement of stomatal index, 10-day-old plants grown under L/D conditions were stained with 50 μg/mL of propidium iodide (PI) and observed using a confocal laser scanning microscope, FV1000 (Olympus) in three square areas of 0.48 mm^2^ per cotyledon from 5 cotyledons of 5 independent plants. The stomatal index was calculated as previously described^24^. Error bars, representing standard errors, were calculated from the results of 5 independent cotyledons.

For root elongation measurements, plants were grown vertically under L/D conditions. Root length was measured at ZT0. For quantification of root meristem size, 7-day-old plants grown under L/D conditions were stained with 20 μg/mL PI and observed using an FV1000 microscope as described above. Meristematic cell numbers were determined from observations of the cortical cells, using confocal microscopy images.

For DNA ploidy analyses, plants immediately before and 28 h after induction in VISUAL were used. Cotyledons were chopped using a razor blade in 0.5 mL of nuclei-extraction buffer (solution A of the Cystain UV precise P; Partec). After filtration through a 30-μm mesh, 2 mL of the staining solution containing DAPI (solution B of the kit) was added. Ploidy levels were measured using a ploidy analyzer PA (Partec). The lowest peak of WT was assumed to represent 2C nuclei (C is the haploid DNA content).

For quantification of cell-cycle progression, S-phase cells were visualized using Click-iT EdU Alexa Fluor 488 Imaging Kits (Thermo Fisher Scientific) according to the manufacturer’s instructions. At various time points, 7-day-old plants grown under L/D conditions were transferred to a plate containing identical medium with 10 μM EdU and incubated for 1 h. Incorporation of EdU was terminated by fixing the plants with 4% paraformaldehyde. EdU incorporated into DNA was stained by Alexa Fluor 488 and observed using FV1000. The numbers of EdU-positive cells in the cortex were counted using ImageJ.

For GUS staining, plants were fixed in 90%(v/v) acetone for 15 min on ice, vacuum-infiltrated and incubated at 37 °C for 2 h (overnight for *IRX3∷GUS*) in the GUS assay solution containing 100 mM sodium phosphate buffer (pH 7.2), 1 mM potassium-ferrocyanide (5 mM for *TDR∷GUS*), 1 mM potassium-ferricyanide (5 mM for *TDR∷GUS*), 0.1%(v/v) Triton X-100 and 0.5 mg ml^−1^ 5-bromo-4-chloro-3-indolyl-β-D-glucuronic acid (X-Gluc). Chlorophylls in the tissue were removed by incubation in 70%(v/v) ethanol.

### Detection of bioluminescence during VISUAL

A cotyledon of a 7-day-old plant grown under L/D conditions was transferred to the liquid medium as described previously^7^, supplemented with 0.025 mM luciferin (Biosynth) at ZT0, and incubated under L/D conditions for 2 days. Then, the cotyledon was transferred to the liquid medium for vascular cell differentiation, containing 0.025 mM luciferin at ZT0 and incubated under LL conditions for 3 days. For photon counting, the emitted luminescence was recorded using a photomultiplier-tube-based bioluminescence monitoring system^25^.

### qRT-PCR

Total RNA was extracted using an RNeasy Plant Mini Kit (QIAGEN) and reverse-transcribed using a Transcriptor First Strand cDNA Synthesis Kit (Roche) according to the manufacturer’s instructions.

Real-time gene expression was analyzed with a CFX96 Real-Time PCR Detection System (Bio-Rad). *UBQ14* were used as an internal control for VISUAL experiments^7^, respectively. Specific sequences for each primer pair were:

CAB3-RT-F, 5′-ACCCAGAGGCTTTCGCGGAGT;
CAB3-RT-R, 5′- TGCGAAGGCCCATGCGTTGT;
TDR-RT-F, 5′- TGGTGGAAGTTACTTTGAAGGAG;
TDR-RT-R, 5′- TTCAATCTCTGTAAACCACCGTAA;
IRX3-RT-F, 5′- CCTCGGCCACAGCGGAGGAT;
IRX3-RT-R, 5′- CGCCTGCCACTCGAACCAGG;
CCA1-RT-F, 5′- GAGGCTTTATGGTAGAGCATGGCA;
CCA1-RT-R, 5′- TCAGCCTCTTTCTCTACCTTGGAGA;
LHY-RT-F, 5′- ACGAAACAGGTAAGTGGCGACA;
LHY-RT-R, 5′- TGCGTGGAAATGCCAAGGGT;
CYCD3;1-qPCR-Fw, 5′- CGAAGAATTCGTCAGGCTCT;
CYCD3;1-qPCR-Rv, 5′- ACTTCCACAACCGGCATATC;
E2Fc-qPCR-Fw, 5′-; GAGTCTCC-CACGGTTTCAG
E2Fc-qPCR-Rv, 5′-; TCACCATCCGGTACTGTTGC
UBQ14-qPCR-Fw, 5′- TCCGGATCAGGAGAGGTT; and
UBQ14-qPCR-Rv, 5′- TCTGGATGTTGTAGTCAGCAAGA.
The following thermal cycling profile was used,
CAB3, 95°C for 10 s, ~40 cycles of 95°C for 10 s, 62°C for 15 s and 72°C for 15 s;
CCA1, 95°C for 60 s, ~40 cycles of 95°C for 10 s, 60°C for 15 s and 72°C for 7 s;
IRX3, 95°C for 60 s, ~40 cycles of 95°C for 10 s, 64.5°C for 15 s and 72°C for 10 s; and
TDR, LHY, CYCD3;1, E2Fc, and UBQ14, 95°C for 60 s, ~40 cycles of 95°C for 10 s, 60°C for 15 s and 72°C for 15 s.

Each sample was run in technical triplicate to reduce experimental errors. Error bars, representing standard errors, were calculated from the results of biological triplicates.

### scRNA-seq and cpRNA-seq

For scRNA-seq, a cotyledon was placed adaxial side down on a glass slide, and fixed with an adhesive tape, e.g., cellophane tape. Then the center of cotyledon was cut using a razor blade, and with the aid of a microscope, the contents of a single cell were collected using a glass capillary. Samples were subjected to UMI-tagged sequencing using a NextSeq 500 system (Illumina). The process closely followed the method described by Kubo *et al*^10^. For cpRNA-seq, total RNA was extracted using an RNeasy Plant Mini Kit and subjected to UMI-tagged sequencing, as for scRNA-seq, except that 10 cycles of the PCR amplification step were required. scRNA-seq data was normalized together with cpRNA-seq by DESeq^26^.

### WISP pipeline

To identify scRNA-seq data related to the xylem cell lineage, Wishbone was performed as previously described^11^. Samples after induction in VISUAL were subjected to Wishbone. The first and third components of principle component analysis (PCA) were used for t-SNE. The xylem lineage was selected according to the expression of xylem cell marker genes on the t-SNE plots.

Clustering of cells was performed using the Seurat R package^12^. In brief, digital gene expression matrices were column-normalized and log-transformed. To obtain a landmark gene set for Seurat, we divided all genes in cpRNA-seq data into 17 groups according to the peak expression time of each gene. The 17 genes showing the highest correlation coefficient with scRNA-seq data in each respective group were selected as landmark genes. In addition to the 17 genes, cell-type-specific markers (*CAB3*, *LHCB2.1*, *TDR*, *AtHB8*, *IRX3*, *IRX8*, and *SEOR1*) were also selected as landmark genes. In total, 24 genes were used as a landmark gene set for Seurat.

Finally, we selected the genes whose correlation coefficient between scRNA-seq and cpRNA-seq was more than 0.5. Among the selected genes, 2,217 genes whose sum total of expression levels in the scRNA-seq data were higher than the value equivalent to ten times the cell numbers were subjected to PeakMatch with the following parameters: T = 2, last = 0, intv = 1, inter = 7.

### Overview of the PeakMatch algorithm

Let *Z* be the set of whole genes under consideration. Discretizing pseudo and actual times into integers for simplicity, we denote by *P* = {1, …, m} and *A* = {1, … , n} the sets of available pseudo and actual times, respectively. Suppose that, for each gene *z* ∈ *Z* , we are given pseudo time-series based scRNA-seq data *S*_*z*_ = (*s*_*z*,1_, … , *s*_*z*,*m*_) and actual time-series based cpRNA-seq data *C*_*z*_ = (*c*_*z*,1_, … , *c*_*z*,*n*_), where *s*_*z*,*p*_ ∈ *S*_*z*_ and *c*_*z*,*a*_ ∈ *C*_*z*_ represent the gene *z*’s expression levels at a pseudo time *p* in the scRNA-seq data, and at an actual time *a* in the cpRNA-seq data, respectively.

To estimate the actual times of gene expressions in the scRNA-seq data, we would like to find pairs (*p*, *a*) ∈ *P* × *A* of pseudo and actual times so that the expression levels *s*_*z*,*p*_ and *c*_*z*,*a*_ are likely to be “comparable” for many genes *z* ∈ *Z*. Once such pairs (*p*, *a*) are found, we may estimate the actual time of *s*_*z*,*p*_ by that of *c*_*z*,*a*_.

The point is that, among the observed gene expression levels, “peaks” are the most important phenomena. Then it is desired that a peak in *S*_*z*_ and a peak in *C*_*z*_ should be matched. It is also required that the pseudo time order should be preserved in the time pairs. To be more precise, whenever a pseudo time *p* is matched to an actual time *a,* any pseudo time *p′* > *p* should be matched to an actual time *a′* > *a*.

We formulated the problem of finding such time pairs as the maximum weighted non-crossing matching (MWNCM) problem for a bipartite graph. The problem is polynomially solvable^27^, meaning that it is efficiently solvable from the standpoint of the theory of computational complexity.

We took the bipartite graph so that one vertex subset was the pseudo time set *P* and the other vertex subset was the actual time set *A*. For the edge set, we considered all possible pairs (*p*, *a*) ∈ *P* × *A*, where we determined the weight of an edge (*p*, *a*) heuristically by how the pseudo time *p* and the actual time *a* were comparable peaks. We determined the weight of an edge (*p*, *a*) as follows. For each gene *z* ∈ *Z*, we decided whether or not the value *s*_*z*,*p*_ (*resp*., *c*_*z*,*a*_) was within a “peak area” in *S*_*z*_ (*resp*., *C*_*z*_). We considered that *s*_*z*,*p*_ (*resp*., *c*_*z*,*a*_) was within a peak area if it was significantly larger than a general trend of *S*_*z*_ = (*s*_*z*,1_, …, *s*_*z*,*m*_) (resp. , *C*_*z*_ = (*c*_*z*,1_, … , *c*_*z*,*n*_)), which was estimated by an exponential moving average. We set the weight of (*p*, *a*) to a larger value if both *s*_*z*,*p*_ and *c*_*z*,*a*_ are among peak areas for more genes.

Given the scRNA-seq and cpRNA-seq data, the algorithm PeakMatch constructed the bipartite graph, derived an MWNCM for it, and estimated the actual times of all pseudo times in *P* based on the derived MWNCM.

For more details and python-based programs, see https://github.com/endo-lab/PeakMatch.

### Plasmid construction

For luciferase reporter constructs, the NOS terminator was amplified by PCR using the following primers:

nosT-F, 5′-CCGCACTCGAGATATCTAGAATCGTTCAAACATTTGGCAA; and
and nosT-R, 5′-TACAAGAAAGCTGGGTCTAGAGATCTAGTAACATAGATGAC.

The amplified fragment was cloned into the XbaI site of pENTR1A (no ccdB) using an In-Fusion HD Cloning Kit (TaKaRa). In the same way, the coding sequence of LUC+ was amplified and cloned into the XhoI-EcoRV site of the plasmid using the following primers:

LUC-XhoI-F, 5′-TTCGCGGCCGCACTCGAGATGGAAGACG; and
LUC-EcoRV-R, 5′-TGAACGATTCTAGATATCTTACACGGCGATCTTTCCGC.

The resulting pENTR1A (LUC-nosT) was used for *LHY∷LUC*. The promoter of *LHY* was amplified from Col-0 genomic DNA using the following primers:

LHY-pro-F (XhoI), 5′-CGCGGCCGCACTCGATTTTGGAATAATTTCGGTTATTTC; and
LHY-pro-R (XhoI), 5′-ATCCGCGGATCTCGAAACAGGACCGGTGCA.

The amplified fragment was cloned into the XhoI site of pENTR1A (LUC-nosT). After sequence verification, the plasmid was recombined with pFAST-G01^28^, introduced into Col-0 plants, and transgenic plants were selected by fluorescence of T1 seeds.

For *IRX3∷GUS*, the promoter of *IRX3* was amplified from Col-0 genomic DNA using the following primers:

IRX3p-Fw (XhoI), 5′-CGCGGCCGCACTCGATCGAGAGCCCGA; and
IRX3p-Rv (XhoI), 5′-GTCTAGATATCTCGAAGGGACGGCCGGAGATTAGCAGCGA.

The amplified fragment was cloned into the XhoI site of pENTR1A (no ccdB). After sequence verification, this plasmid was recombined with pGWB3^29^, and introduced into Col-0 plants.

For *BES1∷BES1-GFP*, the genomic DNA fragment of BES1 containing the 2 kb promoter sequence and the 1 kb downstream sequence from the stop codon was amplified from Col-0 genomic DNA using the following primers:

CACC-pBES1_2k_F, 5′- CACCTCTCAACCTGCTCGGT; and
gBES1-R, 5′- CTCTGATGTGGAGTCAATG.

The amplified fragment was cloned into the pENTR/D-TOPO (Thermo Fisher Scientific). After sequence verification, this plasmid was recombined with pGWB1^29^. The resulting pGWB1-gBES1 was linearized by PCR using the following primers:

gBES1-C_inverse-F, 5′- TGAGATGAAGTATACATGAACCTG; and
bes1_R_nonstop, 5′-ACTATGAGCTTTACCATTTCCAAGCG.

Then, sGFP fragment was amplified using the following primers:

sGFP_for-gBES1-C_F, 5′-GGTAAAGCTCATAGTATGGTGAGCAAGGGCG; and
sGFP_for-gBES1-C_R, 5′-GTATACTTCATCTCACTTGTACAGCTCGTCCATG.

The sGFP fragment was inserted just before the stop codon of BES1 by fusion of the two fragments using In-Fusion HD Cloning Kit. After sequence verification, this plasmid was introduced into the *bes1-3* mutant.

For transactivation assay constructs, the coding sequences of *bes1-D*(L) from cDNA of the *bes1-D* mutant and *RLUC* from the vector pRL vector (Promega) were amplified by PCR using the following primers:

bes1-D(L)-F, 5′-GGACTCTAGAGGATCATGAAAAGATTCTTCTATAATTCC; bes1-D(L)-R, 5′-CGGTACCCGGGGATCTCAACTATGAGCTTTACCATTTCC; RLUC-F, 5′-GGACTCTAGAGGATCATGACTTCGAAAGTTTATGATCC; and RLUC-R, 5′-CGGTACCCGGGGATCTTATTGTTCATTTTTGAGAAC.
The amplified fragments were cloned into the BamHI site of pPZP211/NP/35S-nosT^30^ using an In-Fusion HD Cloning Kit.

### Western blotting

Plants expressing *BES1∷BES1-GFP* were harvested every 8 h from 24 h before induction in VISUAL, up to 72 h after induction. Approximately 50 mg of seedlings were ground into a fine powder in liquid nitrogen with a mortar and pestle, mixed with equal volumes of 2×Laemmli sample buffer (100 mM Tris-HCl(pH 6.8), 4%(w/v) SDS, 10%(v/v) 2-mercaptoethanol, and 20%(v/v) glycerol) and boiled at 95°C for 5 min. Samples were separated by SDS-PAGE on a 7.5% acrylamide gel, and transferred onto polyvinylidene fluoride membranes (Bio-Rad Laboratories). For the primary antibody, polyclonal anti-GFP (MBL-598) was diluted 1:2,000. For the secondary antibody, ECL Rabbit IgG, HRP-linked whole Ab (GE Healthcare) was diluted 1:10,000. Blots were visualized with ECL Prime reagent (GE Healthcare) and ImageQuant LAS 4010 (GE Healthcare).

### ChIP-qPCR and ChIP-seq

Chromatin immunoprecipitation assays using *BES1∷BES1-GFP*/*bes1-3* and *LUX∷LUX-GFP*/*lux-4* were performed as described^31^ with modifications. Briefly, 600 mg of seedlings at 8 h after induction in VISUAL were fixed in PBS containing 1% paraformaldehyde for 10 min at room temperature, and nuclei and chromatin were isolated. The isolated chromatin was sheared with a Covaris S220 sonicator under these parameters: 4–6°C, 175 W peak power, 5% duty factor, 200 cycles/burst, for 50 s of treatment. To immunoprecipitate chromatin, 10 μL of anti-GFP antibody (MBL-598) and 50 μL of Dynabeads Protein G (Thermo Fisher Scientific) were used. The precipitated samples were subjected to qPCR or library preparation for ChIP-seq. MYB30^31^ was used as a positive control for ChIP with BES1. For ChIP-qPCR, specific sequences for each primer pair were:

LHYp-ChIP-1F, 5′- GATTCGGGTAGTTCAGTTCTTCG;
LHYp-ChIP-1R, 5′- GGTTAGTTCGGTTCGGTTCTAGG;
LHYp-ChIP-2F, 5′- CACCGTACCCACTTGTTTAGTCG;
LHYp-ChIP-2R, 5′- CGAGCCAGAAGCTTCAATGTG;
LHYp-ChIP-3F, 5′- GGCTCGTAGAGAAGCAACTTGAG;
LHYp-ChIP-3R, 5′- AGTCATCGCAGATCGACACG;
LHYp-ChIP-4F, 5′- GTGGATTCGTTTGGGTGAGG;
LHYp-ChIP-4R, 5′- AACAGTCGCTGCTTCTCCAG.
MYB30-ChIP-F, 5′-AGGTATTTTACGCTGGAAAATGTGT;
MYB30-ChIP-R, 5′- GAATCATCATAATAAGTATGGAGGTG;
ACT2-ChIP-F, 5′- CGTTTCGCTTTCCTTAGTGTTAGCT; and
ACT2-ChIP-R, 5′- AGCGAACGGATCTAGAGACTCACCTTG.

The following thermal cycling profile was used for all the primers: 95°C for 60 s, ~40 cycles of 95°C for 10 s, 60°C for 45 s.

For ChIP-seq, the sequence libraries were prepared using a TruSeq ChIP Sample Preparation Kit v2 (Illumina), and sequenced using an Illumina NextSeq 500 system with a 75-nt single-end sequencing protocol. The sequence reads were mapped to the TAIR10 *Arabidopsis* genome sequence by HISAT2^33^ with default parameters. Peaks were identified by MACS2^34^, using the matching INPUT control with the genome size parameter “-g 1.3e8”.

### Transactivation assay

*Agrobacterium* cultures carrying plasmids for the transactivation assay were grown overnight at 28°C, collected by centrifugation, and adjusted to an OD600 of 0.4 with infiltration buffer (10 mM MES(pH 5.6), 10 mM MgCl2, 150 μM acetosyringone, and 0.02% Silwet-L77). Cells were kept at 28°C in the dark for 1 to 2 h and then infiltrated into the abaxial air spaces of 4-week-old *N. benthamiana* plants grown under L/D conditions at ZT0. After infiltration, plants were kept under L/D conditions for 36 h and harvested at ZT12. The transactivation assay was performed with a Dual-Luciferase Reporter Assay System (Promega) according to the manufacture’s instructions

### Data Availability

Code for PeakMatch is available online at https://github.com/endo-lab/PeakMatch. Other related, relevant lines and data supporting the findings of this study are available from the corresponding authors upon reasonable request.

Sequence data from this article can be found in The Arabidopsis Information Resource (TAIR) databases (https://www.arabidopsis.org) under the following accession numbers: UBQ14 (At4g02890), ACT2 (At3g18780), LHCB2.1 (At2g05100), CAB2 (At1g29920), CAB3 (At1g29910), AtHB8 (At4g32880), TDR (At5g61480), IRX3 (At5g17420), IRX8 (At5g54690), SEOR1 (At3g01680), COR15A (At2g42540), ADH1 (At1g77120), RD29A (At5g52310), OBP1 (At3g50410), ACS6 (At4g11280), MYB30 (At3g28910), CCA1 (At2g46830), LHY (At1g01060), PRR3 (At5g60100), PRR5 (At5g24470), PRR7 (At5g02810), PRR9 (At2g46790), TOC1 (At5g61380), ELF3 (At2g25930), ELF4 (At2g40080), LUX/PCL1 (At3g46640), BOA/NOX (At5g59570), GI (At1g22770), BES1 (At1g19350), CYCD3;1 (At4g34160), E2Fc (At1g47870), and RBR (At3g12280).

## Acknowledgements

We thank T. Koto, T. Kondo and Y. Sando for technical assistance, J.A. Hejna for English proofreading, N. Mochizuki for the gift of *CAB2∷GUS* seeds, T. Demura for the gift of *TDR∷GUS* seeds, T. Nakano for the gift of *bes1-D* seeds, and N. Takahashi, T. Sugiyama and M. Umeda for advising ploidy experiments. This work was supported by JST PRESTO grant 888067 (to M.E.); by JSPS KAKENHI (grant numbers 15H05958, 16H01240, 17K19392, 18H04781, and 18H02461 (to M.E.), 18K14732 (to K.I.), and 17J08107 (to K.T.)); by ISHIZUE 2017 of the Kyoto University Research Development Program, grants from the Yamada Science Foundation, Senri Life Science Foundation, LOTTE Foundation, Daiichi Sankyo Foundation of Life Science, The Takeda Science Foundation, and the Nakajima Foundation, the Sekisui Chemical Grant Program, SEI Group CSR Foundation, SECOM Science and Technology Foundation, and Tokyo Kasei Chemical Promotion foundation (to M.E.); by Grants-in-Aid for Scientific Research on Priority Area 25113005 (to T.A.). Sample preparation for RNA-seq and ChIP-seq was performed at the Medical Research Support Center, Graduate School of Medicine, Kyoto University.

## Author contributions

K.T., K.I., K.B., H.S., and K.U. performed the gene expression analysis and phenotypic analysis. K.H. and K.T. performed the WISP analysis. T.S., M.K., K.T., H.S., and M.E. performed the single-cell RNA-seq analysis. K.I. performed the ChIP assay. Y.K. assisted with VISUAL experiments. M.S. established *BES1∷BES1-GFP* plants. K.T., K.I., and M.E. wrote the manuscript. M.E. directed and supervised the research with the support of T.A. and H.F..

## Supplementary Information

**Supplementary Table 1.**

Summary of single-cell transcriptome data.

**Supplementary Table 3.**

Reconstructed actual time-series of single-cell transcriptome data using the WISP pipeline.

**Supplementary Table 4.**

Summary of peak calling using the MACS2 and gene annotation of putative LUX target genes.

**Supplementary Video 1. Extracting a single-cell content using a glass capillary.**

**Supplementary Figure 1.**
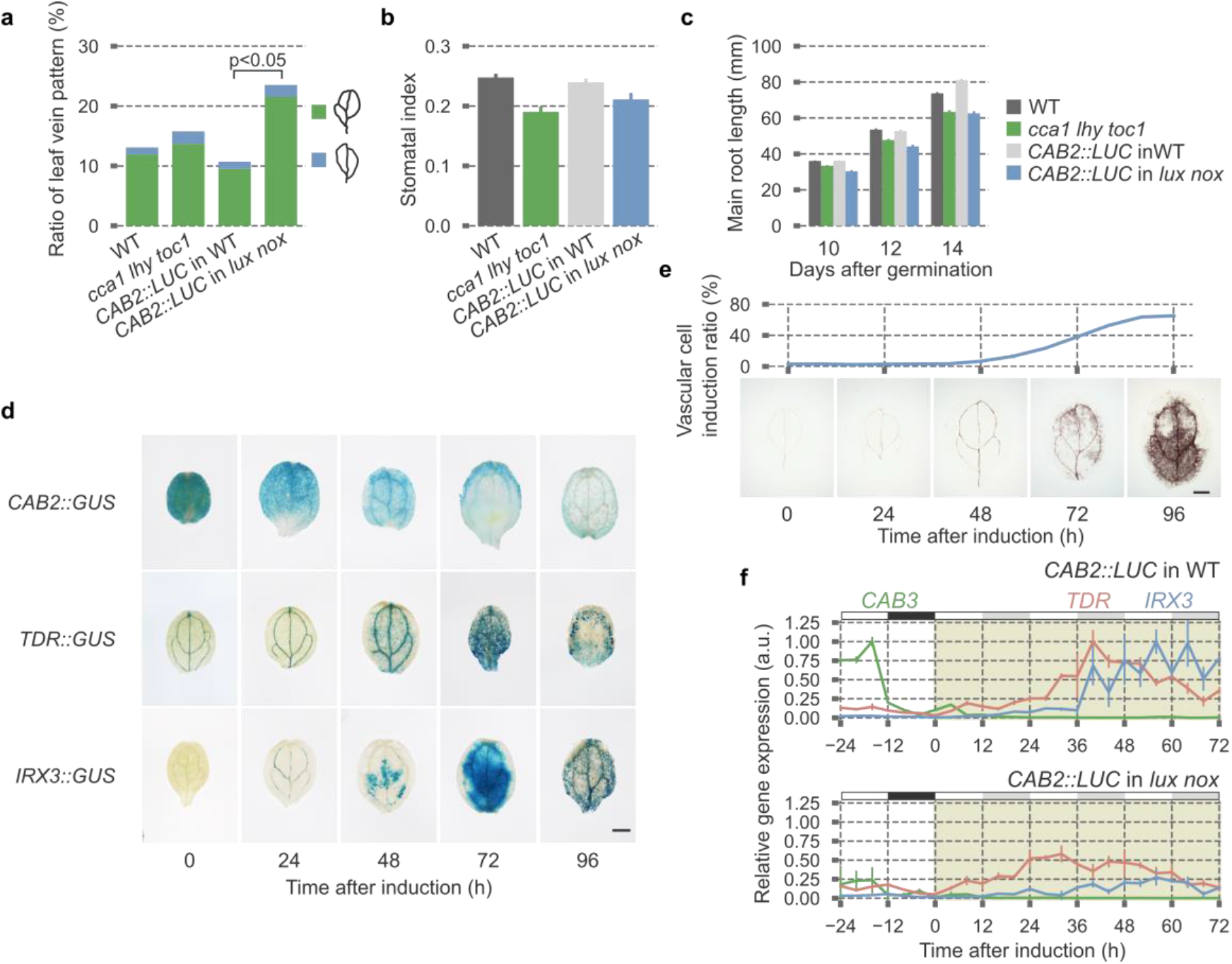
Circadian clock associated with cell differentiation in VISUAL and non-VISUAL. **a**–**c**, Phenotypic analyses of vascular bundle development in cotyledons (**a**, n = 100, chi-square test, p < 0.05), guard cells (**b**, n = 5, mean ± s.e.), and main roots (**c**, n = 10, mean ± s.e.) using WT and clock mutants. Abnormal leaf vein patterns were categorized according to the numbers of areoles (**a**). Stomatal index was calculated as previously described^24^ (**b**). **d**,**e**, GUS staining of cell-type-specific markers (**d**) and the vascular cell induction ratio (**e**) during VISUAL. Representative photos of lignin staining are below (**e**, bar = 1 mm). **f**, Expression patterns of cell-type-specific markers in *lux nox* and corresponding WT during VISUAL (n = 3, mean ± s.e.). Green, red, and blue lines indicate marker genes for mesophyll cells (*CAB3*), vascular stem cells (*TDR*), and xylem cells (*IRX3*), respectively. White, black, and gray boxes indicate light period, night period, and subjective night period, respectively. Peak expression levels of each respective gene in WT were normalized to 1.

**Supplementary Figure 2.**
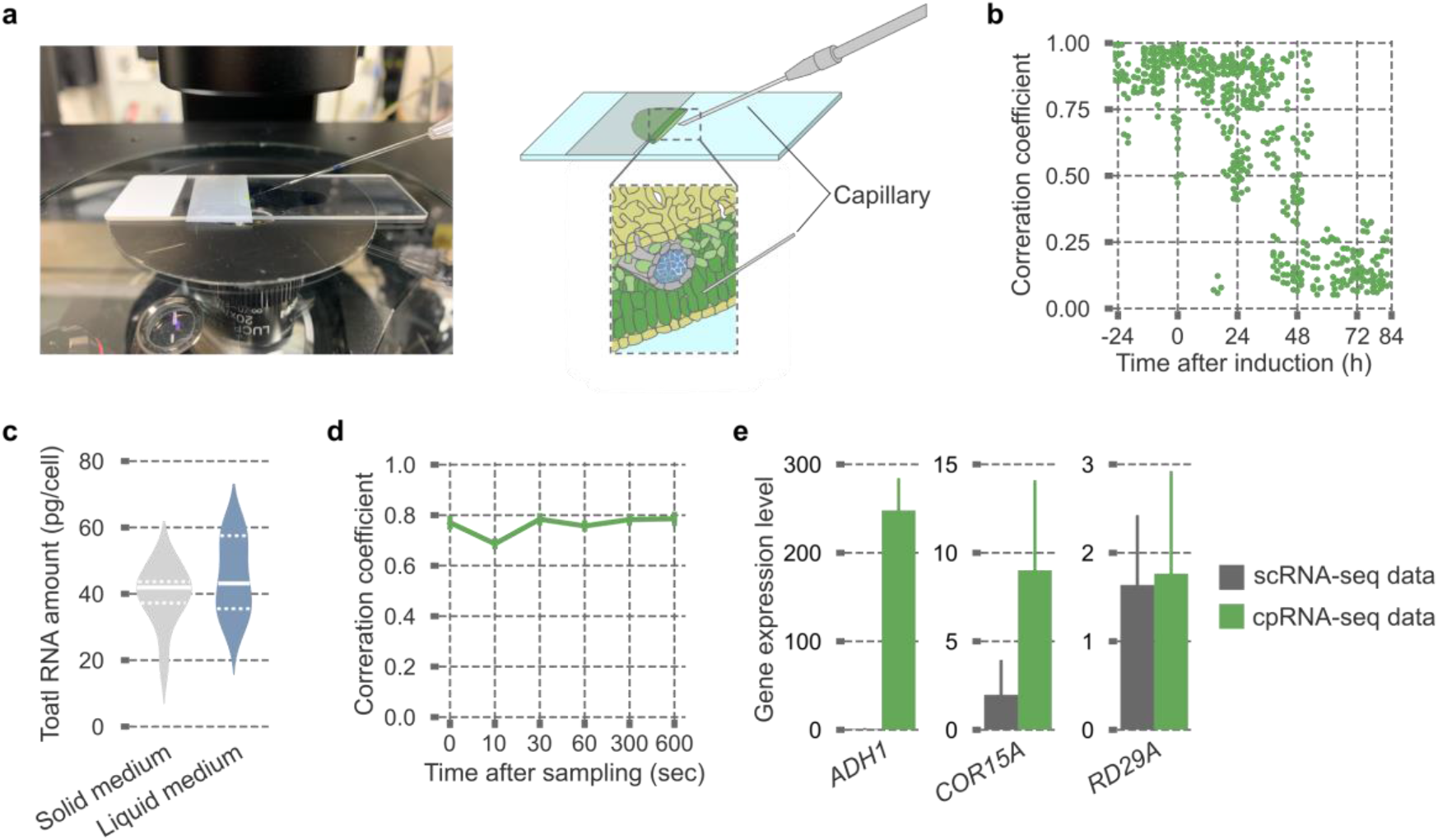
Single-cell transcriptome analysis using microcapillary in *Arabidopsis*. **a**, Set up for microcapillary manipulation to physically extract the contents of individual cells using glass capillaries. **b**, Assessment of scRNA-Seq data during VISUAL. Due to the stochastic cell differentiation in VISUAL, the values of Pearson’s correlation coefficient between samples in the same time points become smaller, suggesting the existence of various types of cells in later time points. **c**, Estimation of total RNA amounts in a single cell. Plants were grown in liquid media and solid agar media, and then isolated cells were analysed as protoplasts. Averaged amounts of total RNA in single cells were comparable (n = 10, mean ± s.e.). White solid lines indicate the median. White broken lines indicate the upper and lower quartiles. **d**, Estimation of mRNA stability during glass capillary-based sampling. In our protocol, harvested single-cell contents were reverse transcribed within 60 s. Even when samples were kept in the glass capillary for 600 s, the values of Pearson’s correlation coefficient between samples from the same time points were almost the same. **e**, Comparison of stress-induced genes expression between cpRNA-seq and scRNA-seq (n = 2 for cpRNA-seq, n = 9 for scRNA-seq, mean ± s.e.). Lower expression levels of *ADH1* and *COR15A* in scRNA-seq indicate that the process of harvesting single cells does not induce stress responses.

**Supplementary Figure 3.**
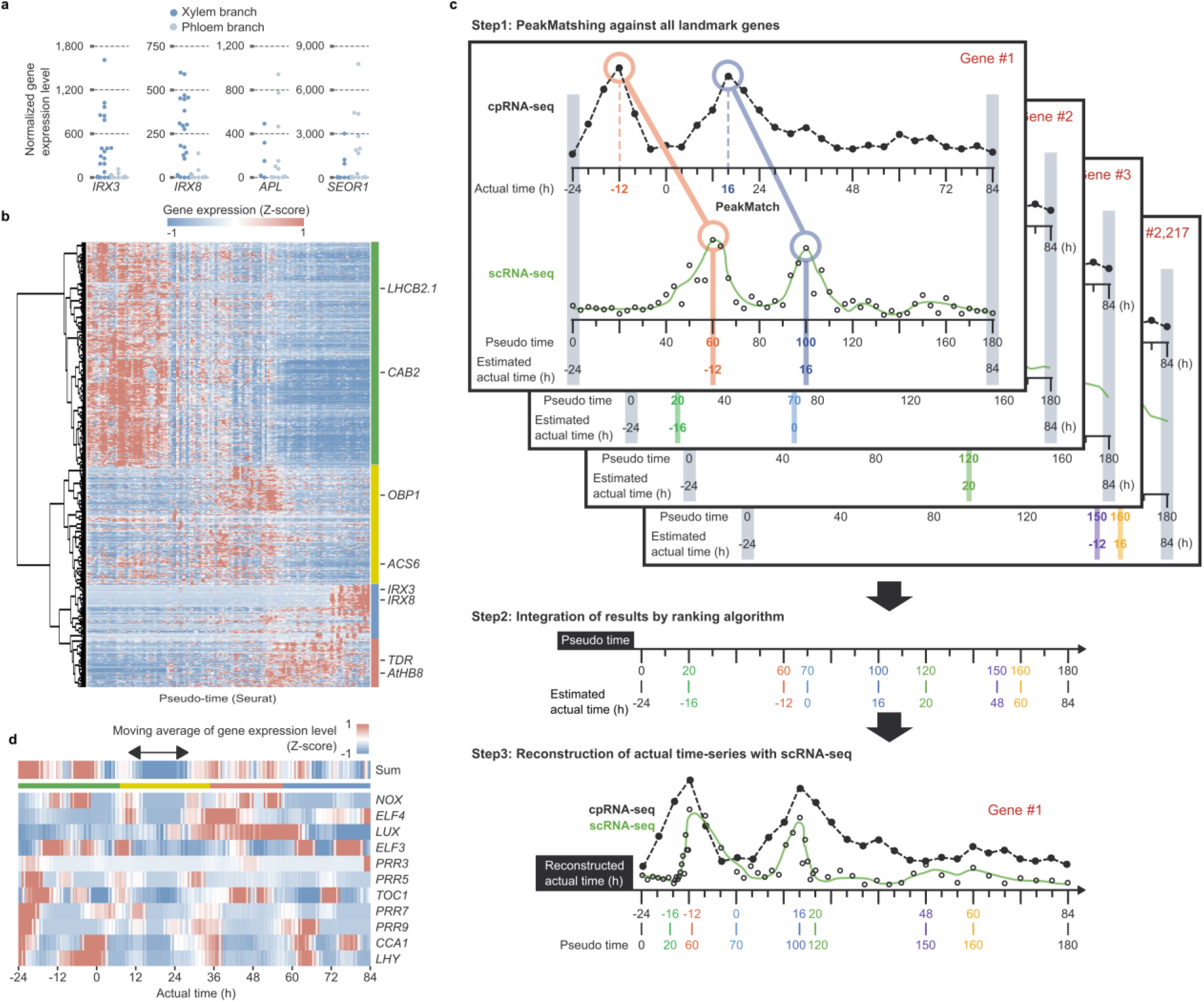
Reconstruction of actual time-series by the WISP pipeline. **a**, Separation of xylem and phloem trajectories by Wishbone. Normalized gene expression levels of marker genes for xylem (*IRX3* and *IRX8*) and phloem (*APL* and *SEOR1*) in each branch are shown. **b**, Hierarchical clustering of pseudo time-series data shows discriminatory gene sets. Green, yellow, red, and blue bars indicate the clusters of mesophyll cell, stem cell, vascular stem cell, and xylem cell states. Marker genes for each cell type are denoted at the right. **c**, A conceptual overview of PeakMatch. The timing of significant gene expression peaks can be comparable between pseudo time-series scRNA-seq data and actual time-series cpRNA-seq data. **d**, A heat map visualizing Z-scores of moving average of each clock gene expression level and their sum total with window size of 4 h. A double-headed arrow indicates a period of time when clock genes cease their rhythmic expression in the stem cell states. In **d** and **e**, green, yellow, red, and blue horizontal bars indicate periods of time corresponding to mesophyll cell, stem cell, vascular stem cell, and xylem cell states, respectively.

**Supplementary Figure 4.**
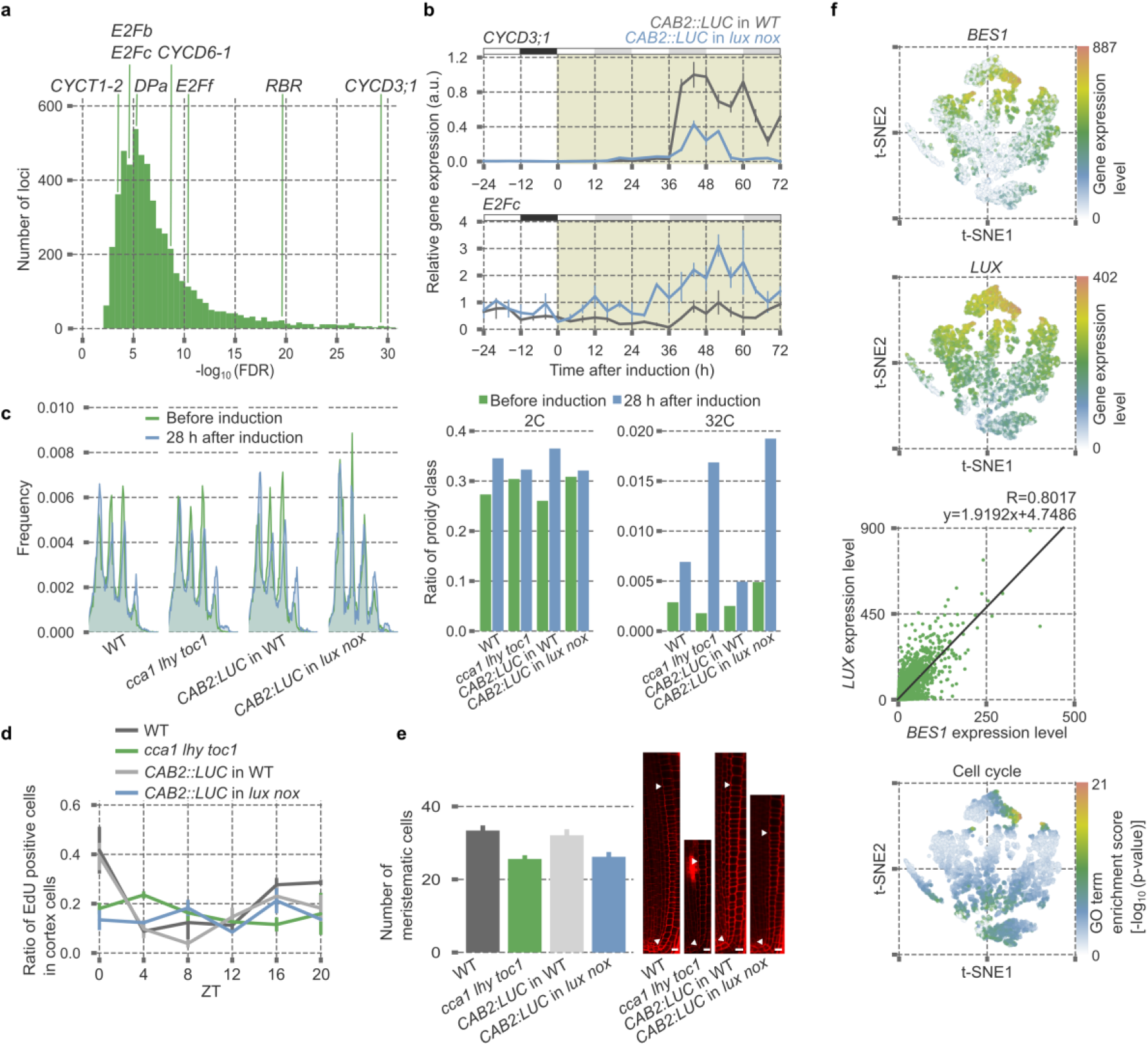
Circadian clock regulates cell-cycle progression. **a**, FDR distributions of genes related to G1-S transition in the candidate LUX target genes. **b**, Expression patterns of *CYCD3;1* and *E2Fc* during VISUAL in *lux nox* and corresponding WT (n = 3, mean ± s.e.). Peak expression levels of each respective gene in WT were normalized to 1. White, black, and gray boxes indicate light period, night period, and subjective night period, respectively. **c**, DNA ploidy analyses before and 28 h after induction using WT and clock mutants (n = 10, mean). Left, DNA ploidy distribution with or without VISUAL. Right, bar graphs of the ratio of 2C and 32C ploidy levels. Clock mutants show increased polyploidy levels probably due to defects in cell-cycle progression. **d**, Incorporation assays of EdU in root meristematic regions using WT and clock mutants under L/D conditions (n = 5, mean ± s.e.). **e**, Comparison of meristematic cell numbers in the root apical meristem using WT and clock mutants (n = 5, mean ± s.e., bar = 10 μm). Representative photos of root meristematic regions are on the right side. Arrowheads indicate the initial and end points of the meristematic zone. **f**, Re-analyses of scRNA-seq data derived from root tissues^20^. Top and second from the top, t-SNE 2D plots overlaid by expression levels of *BES1* and *LUX*. Second from the bottom, correlation coefficient between *BES1* and *LUX* expression levels. Bottom, t-SNE 2D plots overlaid by enrichment scores of the GO-term cell cycle. Genes related to cell cycle are enriched in the cells showing high expression levels of both *BES1* and *LUX*.

**Supplementary Table 2.**
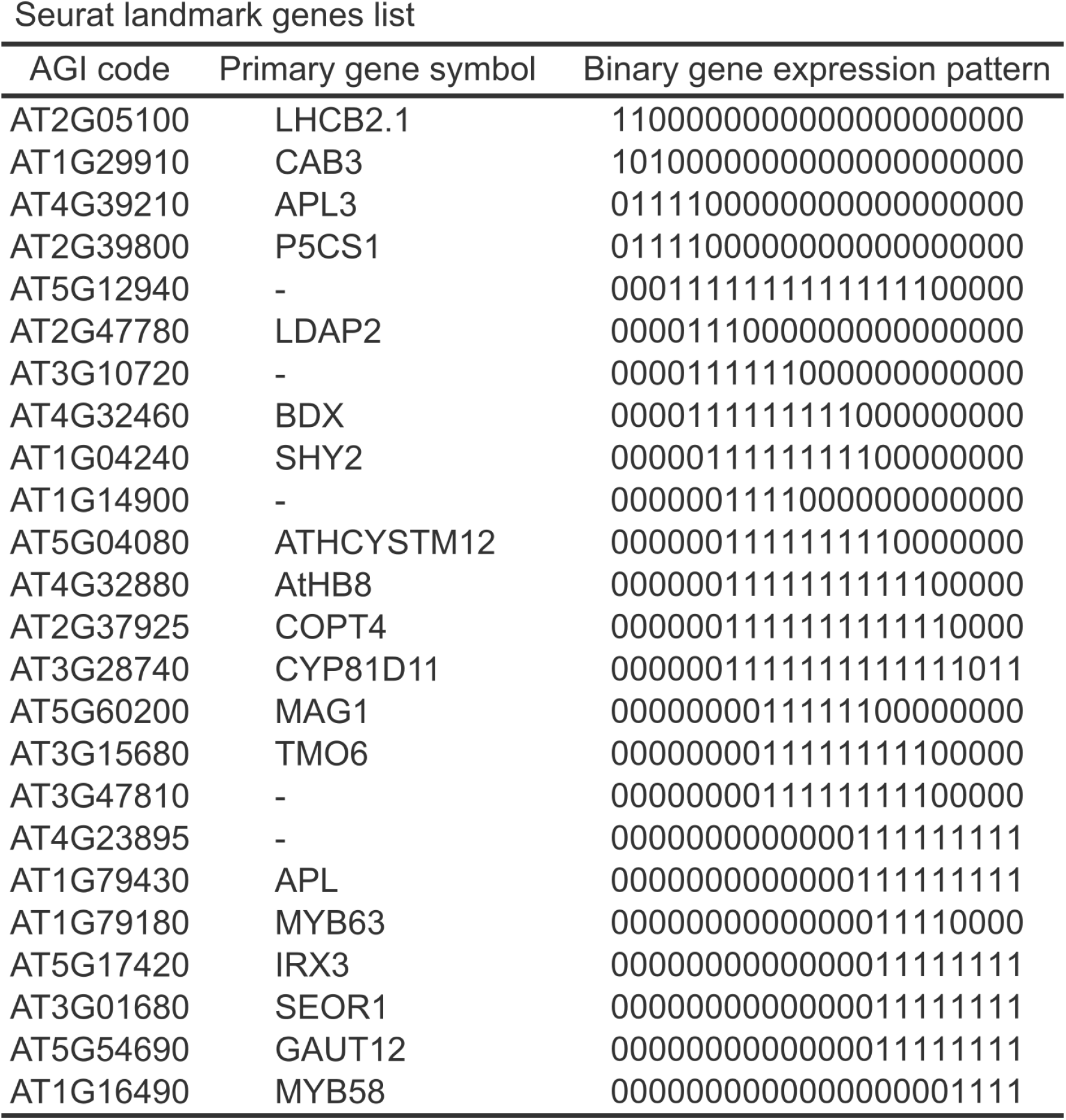
List of a landmark gene set for Seurat

